# Chronically low NMNAT2 expression causes sub-lethal SARM1 activation and altered response to nicotinamide riboside in axons

**DOI:** 10.1101/2024.03.14.584798

**Authors:** Christina Antoniou, Andrea Loreto, Jonathan Gilley, Elisa Merlini, Giuseppe Orsomando, Michael P Coleman

**Affiliations:** John van Geest Centre for Brain Repair, Department of Clinical Neurosciences, University of Cambridge, Forvie Site, Robinson Way, CB2 0PY Cambridge, UK; Department of Clinical Sciences (DISCO), Section of Biochemistry, Polytechnic University of Marche, Via Ranieri 67, Ancona 60131, Italy

**Keywords:** NMNAT2, SARM1, NAD, Programmed axon death

## Abstract

Nicotinamide mononucleotide adenylyltransferase 2 (NMNAT2) is an endogenous axon survival factor that maintains axon health by blocking activation of the downstream pro-degenerative protein SARM1 (sterile alpha and TIR motif containing protein 1). While complete absence of NMNAT2 in mice results in extensive axon truncation and perinatal lethality, the removal of SARM1 completely rescues these phenotypes. Reduced levels of NMNAT2 can be compatible with life, however they compromise axon development and survival. Mice born expressing sub-heterozygous levels of NMNAT2 remain overtly normal into old age but develop axonal defects *in vivo* and *in vitro* as well as behavioural phenotypes. Therefore, it is important to examine the effects of constitutively low NMNAT2 expression on SARM1 activation and disease susceptibility. Here we demonstrate that chronically low NMNAT2 levels reduce prenatal viability in mice in a SARM1-dependent manner and lead to sub-lethal SARM1 activation in morphologically intact axons of superior cervical ganglion (SCG) primary cultures. This is characterised by a depletion in NAD(P) and compromised neurite outgrowth. We also show that chronically low NMNAT2 expression reverses the NAD-enhancing effect of nicotinamide riboside (NR) in axons in a SARM1-dependent manner. These data indicate that low NMNAT2 levels can trigger sub-lethal SARM1 activation which is detectable at the molecular level and could predispose to human axonal disorders.

## Introduction

NMNAT2 (nicotinamide mononucleotide adenylyltransferase 2) is an essential axon survival molecule, the loss of which triggers axon death *in vitro* and *in vivo* [1–3]. Being one of three NMNAT isoforms that catalyse the final step in NAD biosynthesis, NMNAT2 is the predominant enzyme in axons. Mice lacking NMNAT2 die at birth with a severe axonal phenotype, characterised by widespread axon truncation in the peripheral nervous system (PNS) and central nervous system (CNS) [2,3]. Axon loss resulting from NMNAT2 depletion requires the prodegenerative protein and toll-like receptor adaptor SARM1 (sterile alpha and TIR motif containing protein 1). Absence of SARM1 delays degeneration caused by NMNAT2 depletion *in vitro*, and remarkably, completely rescues the axonal outgrowth and perinatal lethality in mice lacking NMNAT2, which remain healthy for up to two years and retain normal innervation of distal leg muscles [4,36].

SARM1 has a critical NAD(P) glycohydrolase (NAD(P)ase) activity. This is regulated by NMN and NAD, which are the substrate and product of NMNATs respectively. NMN activates SARM1 NAD(P)ase by binding to an allosteric site in the autoinhibitory ARM domain [5,6], while NAD competes for binding to the same site and opposes SARM1 activation [6–8]. Loss of labile NMNAT2, the major axonal NMNAT isoform, leads to a rise in axonal NMN and a decline in NAD, resulting in the activation of SARM1 NAD(P)ase and axon degeneration. The interplay between pro-degenerative SARM1 and its pro-survival upstream regulator NMNAT2 is critical for axon degeneration following injury and in several models of neurodegeneration [9,10].

Accumulating evidence supports roles for NMNAT2 loss in human disease. Biallelic loss-of-function (LOF) mutations (R232Q and Q135Pfs*44) in the *NMNAT2* gene have been reported in two stillborn fetuses with fetal akinesia deformation sequence (FADS) and multiple congenital abnormalities [11], resembling the mouse phenotype where complete absence of NMNAT2 leads to perinatal death. Homozygous partial LOF mutations in *NMNAT2* (T94M) were reported in two siblings with childhood onset polyneuropathy and accompanying erythromelalgia that is exacerbated by infection [12]. More recently, two *NMNAT2* missense variants (V98M and R232Q, which confer partial and complete LOF respectively), were identified in two brothers with a progressive neuropathy syndrome, who also have erythromelalgia that worsens with infection [13]. Reduced *NMNAT2* mRNA levels have also been reported in Parkinson’s, Alzheimer’s and Huntington’s disease patients [14,15]. These observations highlight the need to further characterise the mechanisms of NMNAT2-mediated pathogenesis in humans.

Although the interplay between NMNAT2 and SARM1 following complete loss of NMNAT2 is well-established, the effect of chronically low NMNAT2 expression on SARM1 activation is less clear. Crucially, there seems to be widespread variability in *NMNAT2* mRNA levels among individuals in the human population [15], which could underlie differential susceptibility to various neurodegenerative stresses, and the partial LOF coding variants described above may have similar effects. Thus, a better understanding of how low NMNAT2 expression influences SARM1 activation, and in turn axon health, is needed. The remarkable rescue achieved by removing SARM1 in mice lacking NMNAT2 [4,36], raises the intriguing question of whether targeting SARM1 could have such striking outcomes in humans with partial LOF mutations in *NMNAT2* or with low expression level [16].

In addition, NMN and other NAD precursors, such as nicotinamide (NAM) and nicotinamide riboside (NR), are widely used as a strategy to boost NAD levels with the purpose of promoting longevity and healthy aging [17,18]. Initial reports in small cohorts suggest these are safe [19,20] but it is important to ask whether there can be exceptions. In particular, NMN is the endogenous activator of SARM1 and while it does not have adverse effects when NMNAT activity is intact to convert it immediately to NAD and prevent its accumulation, this could differ when NMNAT activity is insufficient. Thus, it is crucial to investigate the impact of these molecules on SARM1 activation and axon integrity under conditions of compromised NMNAT activity, such as in the presence of low NMNAT2 expression in axons [16].

The present study sought to investigate whether constitutively low levels of NMNAT2 can activate SARM1 in morphologically intact axons. We have previously demonstrated that compound heterozygous mice with one silenced and one partially silenced *Nmnat2* allele, which express sub-heterozygous levels of NMNAT2 are overtly normal but present with *in vivo* and *in vitro* axonal defects and behavioural abnormalities [21]. These include an early reduction of myelinated sensory axons with accompanying temperature insensitivity, and a later loss of motor axons with a decline in motor performance. In culture, superior cervical ganglia (SCG) derived from NMNAT2 compound heterozygous mice have impaired neurite outgrowth and are more sensitive to the chemotherapy drug vincristine [21] and the mitochondrial toxin CCCP [22]. However, the underlying mechanism was not previously studied. We now demonstrate that sub-heterozygous NMNAT2 expression in mice reduces the number of live births in a SARM1-dependent manner. Furthermore, chronic and partial SARM1 activation underlies the NAD depletion and neurite outgrowth defect in primary SCG cultures with sub-heterozygous NMNAT2 expression. Most surprisingly, the administration of the NAD precursor NR fails to increase NAD as it does in wild-type or heterozygous cultures, and instead causes a SARM1-dependent depletion of NAD in axons where NMNAT2 levels are low. These data indicate that low NMNAT2 expression leads to sub-lethal SARM1 activation in at least some neuron types, increasing susceptibility to otherwise innocuous stimuli, with potential to prime for axon degeneration disorders in humans.

## Materials and Methods

### Animals

Animal work was approved by the University of Cambridge and performed in accordance with the Home Office Animal Scientific Procedures Act (ASPA), 1986 under project licence P98A03BF9. Animals were kept under standard specific pathogen free (SPF) conditions and fed ad libitum. Mice of both sexes were studied in experiments.

Generation of mice carrying the *Nmnat2^gtE^* and *Nmnat2^gtBay^* gene trap alleles and crosses to generate *Nmnat2^gtBay/gtE^* compound heterozygous mice have been described previously [3,23]. Animals were transferred to a new facility before this work was initiated, which resulted in a bottleneck in numbers. *Nmnat2^gtBay/gtE^* compound heterozygous mice homozygous for the *Sarm1* deletion were generated by crossing *Nmnat2^gtBay/gtE^* mice with *Sarm1* knockout (*Sarm1^-^*

*^/-^*) mice. F1 mice from this cross heterozygous for either *Nmnat2* gene trap allele and heterozygous for the *Sarm1* deletion i.e. *Nmnat2^+/gtBay^;Sarm1^-/+^* and *Nmnat2^+/gtE^;Sarm1^-/+^* were crossed again with *Sarm1* null mice in order to introduce the *Nmnat2* gene trap alleles on a homozygous *Sarm1* null background.

All mice used in this study originated from the same breeding colony and littermates were used wherever possible. While it would be preferable to perform all experiments with equal numbers of *Sarm1* null and *Sarm1* wild-type neurons in parallel, this was usually not possible because the random assortment of genotypes within each litter meant it was neither cost effective nor ethical to breed large enough numbers of mice to enable this.

### Genotyping

Separate duplex polymerase chain reaction (PCR) was performed to assess the presence of each of the two gene trap alleles, *Nmnat2^gtE^* and *Nmnat2^gtBay^*. A duplex PCR was performed to determine the *Sarm1* genotype. Primers 5ʹ-ctcagtcaatcggaggactggcgc-3ʹ (forward for gene trap), 5ʹ-gctggcctaggtggtgatttgc-3ʹ (forward for wild-type) and 5ʹ-cacaaggcctttctcagacttgc-3ʹ (common reverse for both) were used to amplify a 215 bp product from the *Nmnat2^gtE^* gene trap allele and a 389 bp product from the corresponding wild-type locus. Temperatures used were 94 °C for denaturation and 60 °C for primer annealing. Primers 5ʹ-aggaagcagggagaggcag-3ʹ (reverse for wild-type), 5ʹ-tgcaaggcgattaagttgggtaacg-3ʹ (reverse for gene trap) and 5ʹ-gagccacagactagtgactggttg-3ʹ (common forward for both) were used to amplify a 206 bp product from the *Nmnat2^gtBay^* gene trap allele and a 310 bp product from the corresponding wild-type locus. Temperatures used were 94 °C for denaturation and 65 °C for primer annealing. Primers 5ʹ-acgcctggtttcttactctacg-3ʹ and 5ʹ-ccttacctcttgcgggtgatgc-3ʹ were used to amplify a >500 bp product (∼508 bp) from the wild-type *Sarm1* allele and primers 5ʹ-ggtagccggatcaagcgtatgc-3ʹ and 5ʹ-ctcatctccgggcctttcgacc-3ʹ were used to amplify a <500 bp product (∼450 bp) from the neomycin resistance cassette retained in the knockout allele in place of deleted exons 3-6 [24]. Temperatures used were 94 °C for denaturation and 60 °C for primer annealing.

### Primary neuronal explant cultures

Superior cervical ganglia (SCGs) were dissected from P0-P3 mouse pups and dorsal root ganglia (DRGs) were dissected from E13-14 mouse embryos. Explants were plated in 3.5 cm tissue culture dishes pre-coated with poly-L-lysine (20 mg/ml for 1 hour; Sigma) and laminin (20 mg/ml for 1–2 hours; Sigma). Explants were cultured in Dulbecco’s Modified Eagle’s Medium (DMEM) with high glucose, glutamine and sodium pyruvate (Gibco), with 1% penicillin/streptomycin (Invitrogen), 50 ng/ml 2.5S NGF (Invitrogen), and 2% B-27 (Gibco). Aphidicolin 4 μM (Calbiochem) was used to restrict the proliferation and viability of small numbers of non-neuronal cells.

Nicotinamide riboside (NR) was prepared as a 100 mM stock from Tru Niagen capsules (Chromadex) and stored at 4 °C. The contents of the capsules were dissolved in PBS without Ca^2+^ and Mg^2+^ (Merck) and passed through a 0.22 μm filter. Nicotinamide (NAM) (Sigma-Aldrich) was prepared in water and stored frozen as 100 mM stock aliquots. Cell culture media was supplemented with NR (2 mM) or NAM (1 mM) at the days *in vitro* (DIV) indicated in the figures.

### Acquisition of neurite images and quantification of neurite outgrowth

Phase contrast images were acquired on a DMi8 upright fluorescence microscope (Leica microsystems) coupled to a monochrome digital camera (Hammamatsu C4742-374 95). Neurite outgrowth from SCG and DRG explants was assessed from low magnification (NPLAN 5x/0.12 objective) images on the DIV indicated in the figures. Two measurements of radial outgrowth were recorded for each ganglion, by taking the maximal outgrowth from the edge of the ganglion to the point where the bulk of neurites terminated. The average length was calculated for each day.

### Confocal imaging of PAD6 in primary neuronal cultures

Compound PC6, synthesised and provided by AstraZeneca, was used to visualise SARM1 activation in primary neuronal cultures through its conversion to PAD6 [29]. PC6 was administered in cell culture media at 50 μM and images were acquired 30 minutes later using a Confocal-LSM780 (1.4a) microscope, 40x oil objective, Ex/Em: 405/525 nm. Two representative images were taken for each sample.

### Measurement of NAD and NADP levels from primary neuronal cultures

NAD and NADP levels were measured using the commercially available kits NAD/NADH-Glo Assay and NADP/NADPH-Glo Assay (Promega G9071, Promega G9081), respectively. Neurons were collected in Eppendorf tubes by disrupting adhesion to the plate with a jet of medium and washed twice in ice-cold PBS with Ca^2+^ and Mg^2+^ (Merck), supplemented with complete, ethylenediaminetetraacetic acid (EDTA)-free protease inhibitor cocktail tablets (plus protease inhibitors) (Roche). For experiments where separate measurements were made in ganglia versus neurites, a scalpel was used to cut around and isolate the ganglia and the two compartments were collected in separate tubes. Neurons were lysed in ice-cold Pierce IP lysis buffer (Sigma) plus protease inhibitors. Lysates were centrifuged for 10 min at 13k rpm in a microfuge at 4 °C to pellet insoluble material. Supernatants were collected on ice and diluted to 0.15 μg/μl in ice-cold Pierce IP lysis buffer plus protease inhibitors after protein concentrations had been determined using the Pierce BCA assay (Thermo Fisher Scientific). For NAD and NADP measurements, 25 μl of each extract was mixed with 12.5 μl 0.4 M HCl and heated to 60 °C for 15 min before being allowed to cool at room temperature (RT) for 10 minutes. Reactions were neutralised by adding 12.5 μl 0.5 M Tris base and 10 μl of each neutralised reaction was mixed with 10 μl of the NAD-Glo or NADP-Glo reagent (prepared following manufacturer’s instructions) on ice in wells of a 384-well white polystyrene microplate (Corning). The plate was incubated for 40 min at RT before reading luminescence using GloMax Explorer (Promega) plate reader. Concentrations of NAD and NADP were determined from standard curves generated from dilution series of the relevant nucleotides. Values are expressed as nmol/mg of protein.

### Immunoblotting

Neuronal cultures were collected in Eppendorf tubes by disrupting adhesion to the plate with a jet of medium and washed twice in ice-cold PBS with Ca^2+^ and Mg^2+^ plus protease inhibitors. Neurons were directly lysed into 15 μl 2x Laemmli buffer containing 10% 2-mercaptoethanol and samples were incubated at 100 °C for 5 min. The total amount (15 μl) for each sample was loaded on a 4-20% SDS-PAGE (Bio-Rad). For comparing protein levels between SCG and DRG cultures, samples were first diluted to the same protein concentration, which was determined using the Pierce BCA assay. Briefly, following the washes in PBS, neurons were lysed in ice-cold Pierce IP lysis buffer plus protease inhibitors. Lysates were centrifuged for 10 min at 13k rpm in a microfuge at 4 °C to pellet insoluble material. Supernatants were collected on ice and diluted to the same concentration (based on the sample with the lowest concentration) in ice-cold Pierce IP lysis buffer. Samples were diluted 1 in 2 with 2 x Laemmli buffer and were incubated at 100 °C for 5 min. For detecting NAMPT, SARM1 and GAPDH, 1/6 of the total amount was loaded on a 4-20% SDS-PAGE (Bio-Rad), while the remaining sample was used for detecting NMNAT2. Samples were transferred to Immobilon-FL PVDF membrane (Millipore) using the BioRad Mini-PROTEAN III wet transfer system. Blots were blocked in Tris buffered saline (TBS) (20 mM Tris p.H. 8.3, 150 mM NaCl) with 0.05% Tween 20 (Merck) (TBST) and 5% skimmed milk powder, for 1 hour at RT. Blots were incubated overnight at 4 °C with primary antibodies in TBST containing 5% milk. After three × 10 min washes in TBST, blots were incubated for 1 hour at RT with appropriate HRP-conjugated secondary antibodies (Bio-Rad) diluted 1 in 3,000 in TBST with 5% milk. After three × 10 min washes in TBST blots were incubated with Pierce ECL Western Blotting Substrate or SuperSignal West Dura Extended Duration Substrate (both Thermo Fisher Scientific) and imaged using an Alliance chemiluminescence imaging system (UVITEC Cambridge). Fiji software was used to determine the relative intensities of specific bands from captured digital images.

The following primary antibodies were used: mouse anti-SARM1 monoclonal antibody (1 in 5000, [25]), mouse anti-NAMPT monoclonal antibody (1 in 2000, Cayman Chemical 10813), mouse anti-NMNAT2 monoclonal antibody (1 in 250, Merck WH0023057M1), mouse anti-GAPDH monoclonal antibody (1 in 2000, Abcam ab8245).

### Statistical Analysis

Statistical analysis was conducted using Prism Software (GraphPad Software Inc, La Jolla, CA, USA). The n numbers and specific statistical tests used for each experiment are described in the figure legends. A p value < 0.05 was considered significant (*p < 0.05; **p < 0.01; ***p < 0.001; ****p < 0.0001).

## Results

### Sub-heterozygous NMNAT2 expression reduces viability in a SARM1-dependent manner

In an attempt to investigate the effects of sub-heterozygous levels of NMNAT2 expression, mice heterozygous for two distinct *Nmnat2* gene trap alleles were crossed. Although a gene trap cassette is located in the first intron of the *Nmnat2* gene in each case, the degrees of gene silencing differ, with the *Nmnat2^gtE^* allele being completely silenced and the *Nmnat2^gtBay^* allele being only partially silenced [3,21]. We have previously reported that mice of all four genotypes generated from these crosses are born quite close to the expected ratios [21], although *Nmnat2^gtBay/gtE^* mice, which express sub-heterozygous levels of NMNAT2, were slightly under-represented relative to the expected frequencies without this effect reaching statistical significance. However, we now report a significant loss in prenatal viability of *Nmnat2^gtBay/gtE^* mice, as well as a similar trend to lower than expected numbers at embryonic stage E13-E14 (which may only fail to reach statistical significance because of the smaller sample size) (fig. 1a, b; supplementary fig. 1). Remarkably, knocking out *Sarm1* rescues this prenatal loss, restoring the *Nmnat2^gtBay/gtE^* genotype ratio to the expected level (fig. 1c). Thus, sub-heterozygous NMNAT2 expression can lead to a modest but significant SARM1-dependent loss of viability that is likely to manifest at an embryonic stage. Potential explanations for why one of these studies crosses the threshold for significance and the other did not are discussed below.

**Fig. 1.**
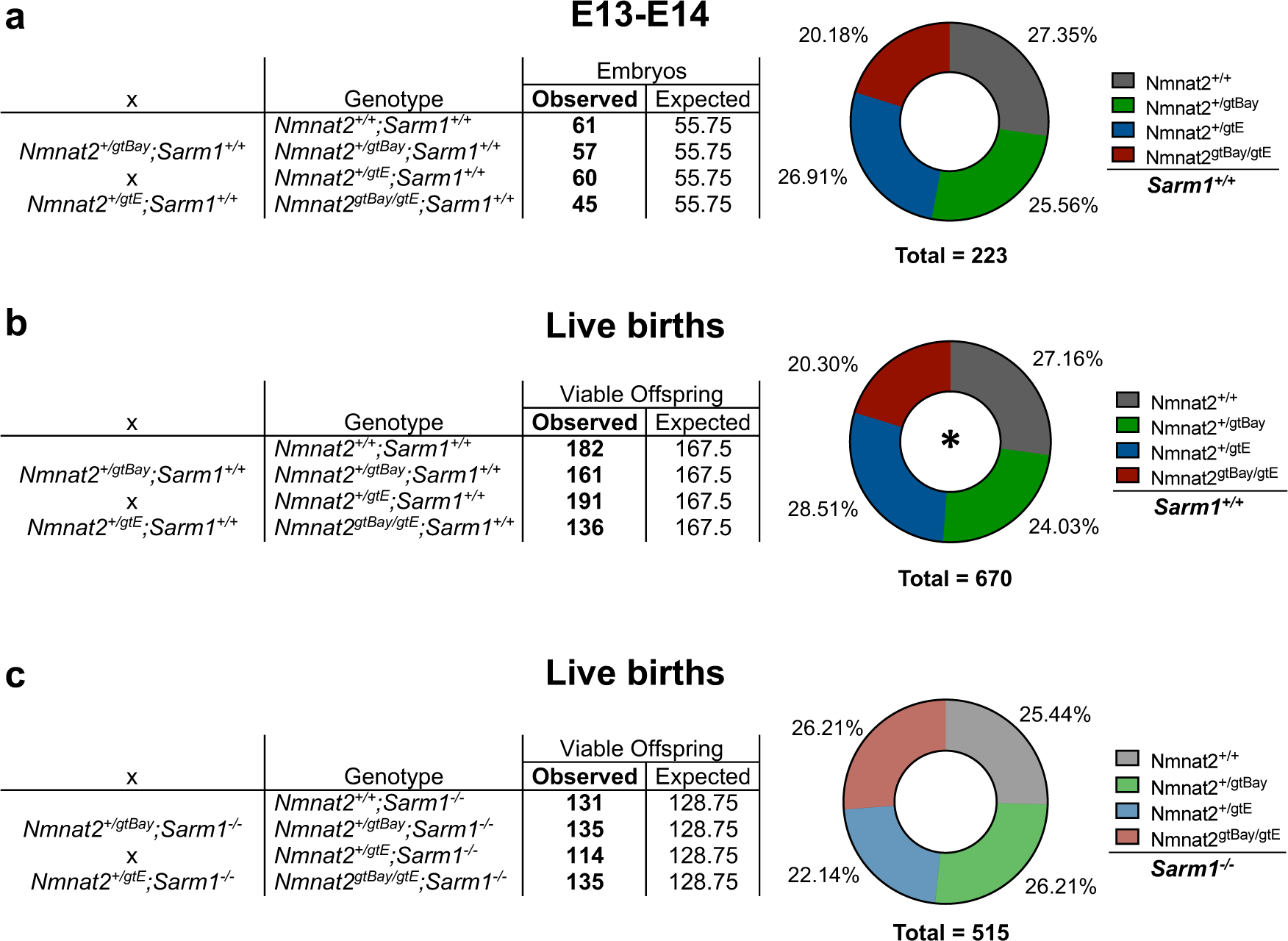
Absence of SARM1 rescues prenatal lethality in *Nmnat2^gtBay/gtE^* mice. (**a**) Genotype frequencies of embryos from crosses between *Nmnat2^+/gtE^* and *Nmnat2^+/gtBay^* mice on a *Sarm1^+/+^* background. The observed embryo frequencies are not significantly different from expected frequencies: *χ*^2^ = 2.919, d.f. = 3, p = 0.4042. (**b**) Genotype frequencies of viable offspring from crosses between *Nmnat2^+/gtE^* and *Nmnat2^+/gtBay^* mice, on a *Sarm1^+/+^* background. The observed birth frequencies are significantly different from expected frequencies: *χ*^2^ = 10.728, d.f. = 3, p = 0.0133. (**c**) Genotype frequencies of viable offspring from crosses between *Nmnat2^+/gtE^* and *Nmnat2^+/gtBay^* mice, on a *Sarm1^-/-^* background. The observed birth frequencies are not significantly different from expected frequencies: *χ*^2^ = 2.336, d.f. = 3, p = 0.5057. Viable offspring numbers include animals between P0-P3 and post-weaning.

### Sub-lethal SARM1 activation underlies the NAD(P) decrease and neurite outgrowth defect in SCG neurons from Nmnat2^gtBay/gtE^ mice

Injury and other insults that activate SARM1 induce its enzymatic activity resulting in the consumption of NAD prior to degeneration [26]. As well as NAD-consuming activity, SARM1 is also an NADPase, cleaving the phosphorylated form of NAD, NADP [27,28]. In an attempt to test for molecular markers of SARM1 activation in non-degenerating axons, NAD and NADP levels were measured in SCG whole explant cultures from wild-type (*Nmnat2^+/+^*), *Nmnat2^+/gtE^* and *Nmnat2^gtBay/gtE^* mice. While NAD and NADP levels in SCG neurons from *Nmnat2^+/+^* and *Nmnat2^+/gtE^* mice were indistinguishable, neurons from *Nmnat2* compound heterozygotes had significantly lower levels of both metabolites (fig. 2a). NAD levels were ∼50% lower in *Nmnat2^gtBay/gtE^* neurons, while a ∼20% reduction in NADP levels was also observed. Thus, halving of NMNAT2 levels does not lower NAD or NADP, whereas sub-heterozygous NMNAT2 expression significantly reduces the levels of both metabolites in SCG primary cultures.

**Fig. 2.**
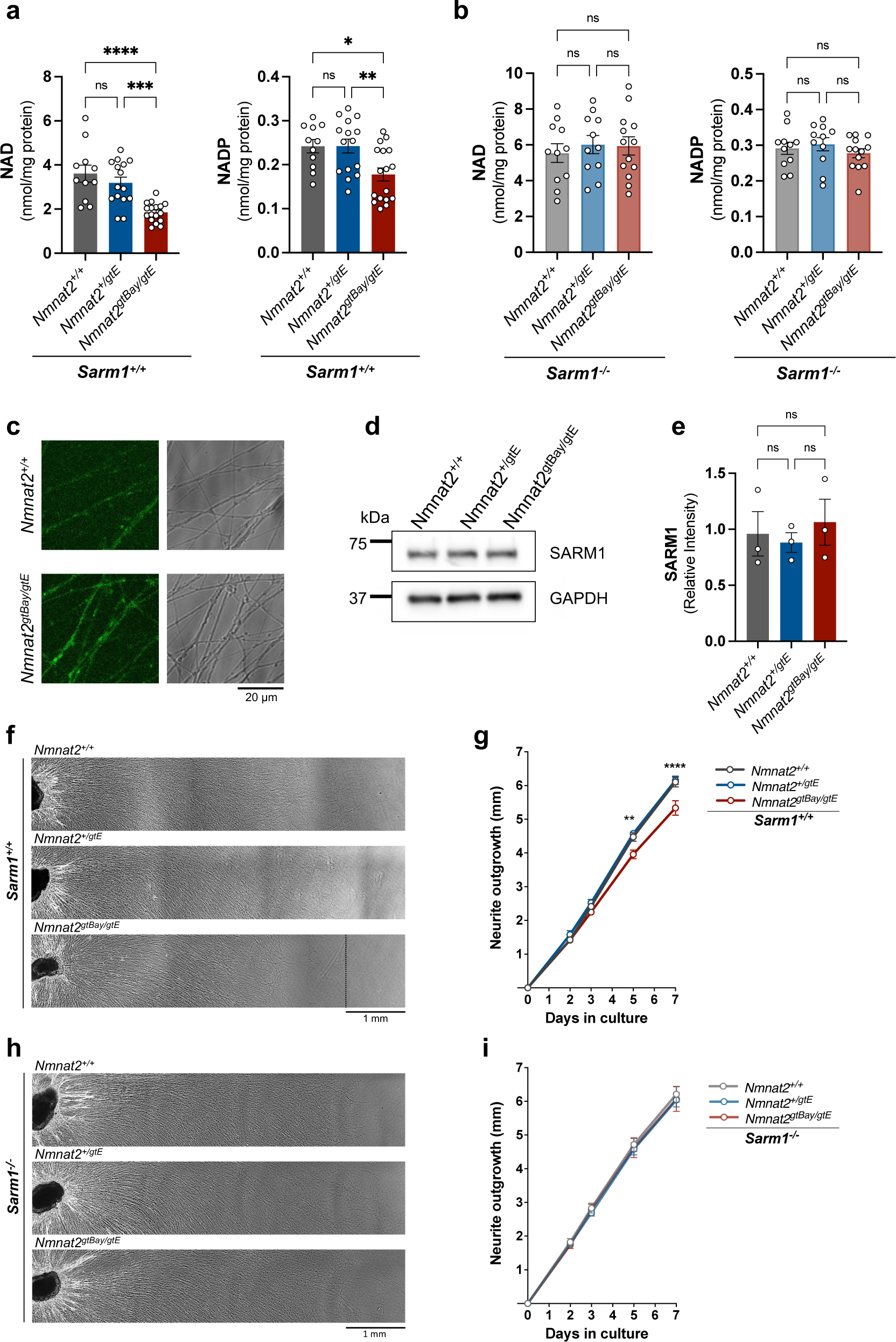
Absence of SARM1 restores NAD(P) levels and rescues the neurite outgrowth defect in SCG neurons from *Nmnat2^gtBay/gtE^* mice. (**a**) NAD and NADP levels in SCG explants of the indicated genotypes on a *Sarm1^+/+^* background (mean ± SEM; n = 11-17 pups per genotype; ****p < 0.0001, ***p < 0.001, **p < 0.01, *p < 0.05 and ns (not significant) = p > 0.05, one-way ANOVA with Tukey’s multiple comparisons test). (**b**) NAD and NADP levels in SCG explants of the indicated genotypes, all on a *Sarm1^-/-^* background (mean ± SEM; n = 11-13 pups per genotype; ns (not significant) = p > 0.05, one-way ANOVA with Tukey’s multiple comparisons test). For a-b, cultures were collected at DIV7. (**c**) Representative images (from n=3) of *Nmnat2^+/+^* and *Nmnat2^gtBay/gtE^* SCG neurons (on a *Sarm1*^+/+^ background) 30 min after incubation with PC6 (50 μM). (**d**) Representative immunoblot of SCG neurite extracts of the indicated genotypes (Sarm1^+/+^) probed for SARM1 and GAPDH (loading control). Cultures were collected at DIV7. (**e**) Quantification of normalised SARM1 levels (to GAPDH) in SCG neurite extracts for the indicated genotypes (*Sarm1*^+/+^) (mean ± SEM; n = 3; ns (not significant) = p > 0.05, one-way ANOVA with Tukey’s multiple comparisons test). (**f**) Representative images of neurite outgrowth at DIV7 in SCG explant cultures of the indicated genotypes on a *Sarm1^+/+^* background. (**g**) Quantification of neurite outgrowth in SCG explant cultures of the indicated genotypes on a *Sarm1^+/+^* background, between DIV0 and DIV7 (mean ± SEM; n = 8-9 pups per genotype; ****p < 0.0001 and **p < 0.01, two-way repeated measures ANOVA with Tukey’s multiple comparisons test for between genotype effects at each time point. Significance is shown for the *Nmnat2^+/+^* vs *Nmnat2^gtBay/gtE^* comparison). (**h**) Representative images of neurite outgrowth at DIV7 in SCG explant cultures of the indicated genotypes on a *Sarm1^-/-^* background. (**i**) Quantification of neurite outgrowth in SCG explant cultures of the indicated genotypes on a *Sarm1^-/-^* background, between DIV0 and DIV7 (mean ± SEM; n = 7 pups per genotype; ns (not significant) = p > 0.05, two-way repeated measures ANOVA with Tukey’s multiple comparisons test for between genotype effects at each time point).

NMNAT2 is an NAD-synthesising enzyme. It is therefore possible that the depletion of NAD(P) observed in primary cultures of mice with sub-heterozygous NMNAT2 expression reflects only a lack of NAD synthesis (due to limited NMNAT2) rather than increased consumption of NAD (due to activated SARM1). In order to establish whether the observed NAD and NADP depletion are SARM1-dependent, *Nmnat2^gtBay/gtE^* mice were crossed to the *Sarm1^-/-^* mice to introduce the *Sarm1* deletion to the *Nmnat2* compound heterozygote mice. Remarkably, absence of SARM1 completely rescues the NAD and NADP depletion in SCG neurons from low-NMNAT2 expressing mice (fig. 2b), providing evidence in support of increased SARM1 activity in morphologically intact, non-degenerating axons.

As an independent, more direct indication of SARM1 activation in *Nmnat2^gtBay/gtE^* SCG neurites, PC6, a recently developed marker of SARM1 activation was used. PC6 is a pyridine base that gets converted by SARM1-dependent base exchange to PAD6, a molecule with increased fluorescence emission at 525nm [29]. Incubation of primary SCG neurons with PC6 gave a markedly higher signal in primary SCG neuron cultures from *Nmnat2^gtBay/gtE^* mice compared to wild-type controls (fig. 2c), suggesting that SARM1 is chronically activated in these axons. No significant difference in SARM1 protein levels was observed between *Nmnat2^+/+^* and *Nmnat2^gtBay/gtE^* SCG neurons (fig. 2d, e), ruling out the possibility that the difference in the PAD6 signal is due to variability in SARM1 levels between the two genotypes.

We previously demonstrated that complete absence of NMNAT2 severely compromises neurite outgrowth in primary neuronal cultures, whereas a 50% reduction in NMNAT2 levels has no detectable phenotype [3]. Interestingly, sub-heterozygous levels of NMNAT2 are consistent with an intermediate phenotype, as SCG neurites from *Nmnat2^gtBay/gtE^* mice have a reduced outgrowth rate compared to both wild-types and *Nmnat2^+/gtE^* single heterozygotes, especially once neurites extend beyond several millimetres [21]. Here we show that absence of SARM1 restores outgrowth of *Nmnat2^gtBay/gtE^* neurites to control levels (fig. 2f-i), demonstrating that the defect is SARM1-dependent and is not solely driven by the lower levels of NMNAT2 expression.

### Sub-heterozygous NMNAT2 expression does not result in NAD(P) depletion or neurite outgrowth defect in DRG neurons

Having obtained evidence in favour of sub-lethal SARM1 activation in SCG neurons, we next addressed whether low NMNAT2 expression has a similar effect in other neuron types. DRG neurons from E13-E14 embryos of the same *Nmnat2* genotypes (i.e. *Nmnat2^+/+^*, *Nmnat2^+/gtE^, Nmnat2^gtBay/gtE^*) were used. Surprisingly, in contrast to our findings in SCG cultures, no NAD or NADP depletion was observed in DRG neurons from *Nmnat2^gtBay/gtE^* mice (fig. 3a). The fact that nucleotide measurements were made in whole explant cultures, encompassing both the cell body and neurite compartments, prompted us to ask whether the presence of NMNAT1, the nuclear NAD-synthesising enzyme, could mask a potential NAD loss occurring solely in the axons where the effect of NMNAT2 is more prominent. For this reason, metabolite measurements were made separately in cell bodies versus neurites. However, no significant difference in NAD or NADP was observed among the three genotypes in separate cell body or neurite fractions (fig. 3b, 3c), although a downwards trend with lower NMNAT2 expression was observed in the neurites (fig. 3c). In addition, in agreement with our previous report [21], we found no evidence for any outgrowth defect in *Nmnat2^gtBay/gtE^* DRG cultures (fig. 3d, e).

**Fig. 3.**
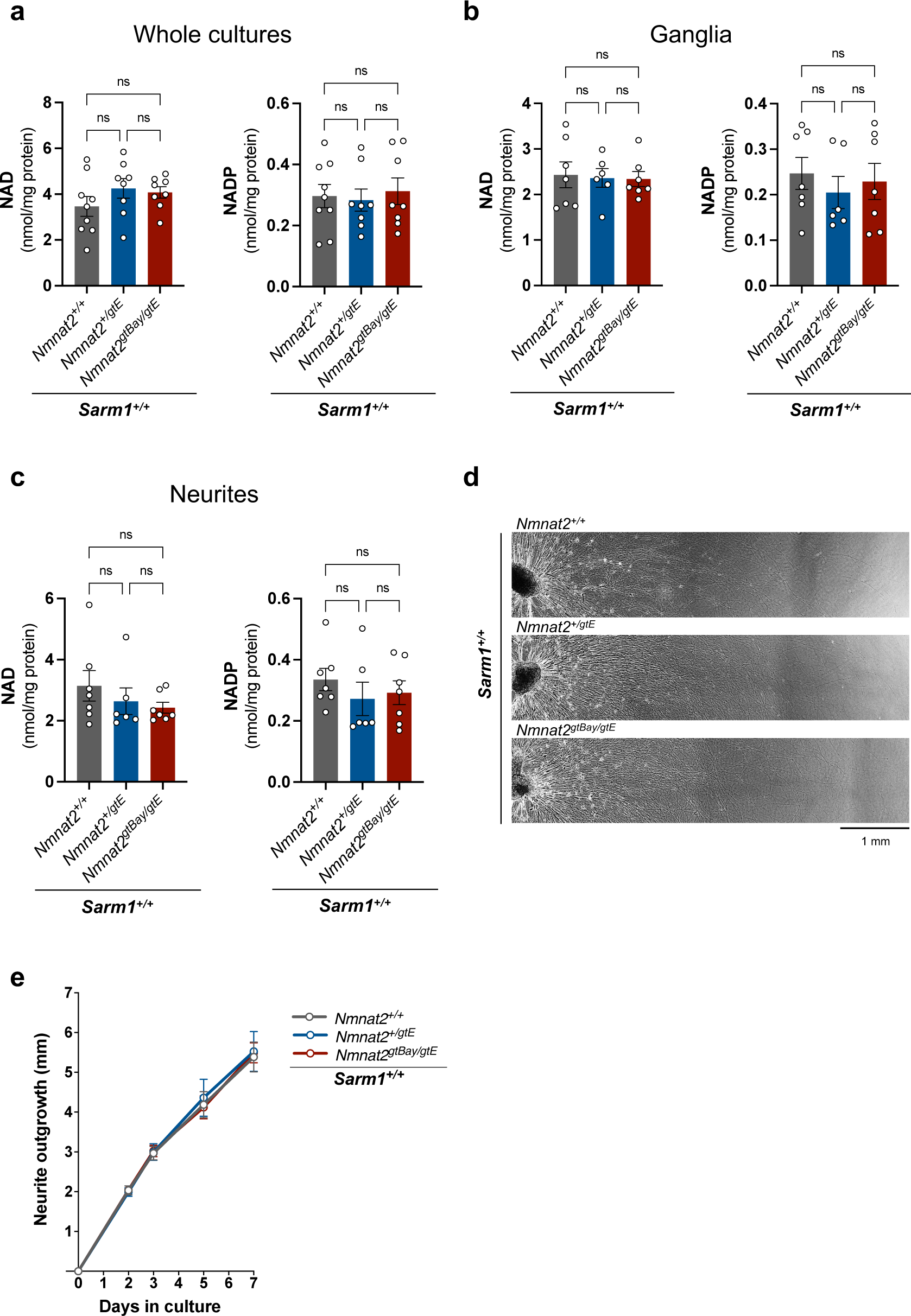
No NAD(P) depletion or neurite outgrowth defect in DRG neurons from *Nmnat2^gtBay/gtE^* mice. (**a**) NAD and NADP levels in DRG whole explant cultures of the indicated genotypes on a *Sarm1^+/+^* background (mean ± SEM; n = 8-9 embryos per genotype; ns (not significant) = p > 0.05, one-way ANOVA with Tukey’s multiple comparisons test). (**b**) NAD and NADP levels in DRG ganglia of the indicated genotypes on a *Sarm1^+/+^* background (mean ± SEM; n = 6-7 embryos per genotype; ns (not significant) = p > 0.05, one-way ANOVA with Tukey’s multiple comparisons test). (**c**) NAD and NADP levels in DRG neurites of the indicated genotypes on a *Sarm1^+/+^* background (mean ± SEM; n = 6-7 embryos per genotype; ns (not significant) = p > 0.05, one-way ANOVA with Tukey’s multiple comparisons test). For a-c cultures were collected at DIV7. (**d**) Representative images of neurite outgrowth at DIV7 in DRG explant cultures of the indicated genotypes on a *Sarm1^+/+^* background. (**e**). Quantification of neurite outgrowth in DRG explant cultures of the indicated genotypes on a *Sarm1^+/+^* background, between DIV0 and DIV7 (mean ± SEM; n = 3-4 embryos per genotype; ns (not significant) = p > 0.05, two-way repeated measures ANOVA with Tukey’s multiple comparisons test for between genotype effects at each time point).

Our observations indicate that different neuron types display varying susceptibility to NMNAT2 depletion. In an attempt to identify possible mechanisms rendering SCG neurons more susceptible, the expression levels of proteins involved in NAD synthesis (NAMPT, NMNAT2) and NAD consumption (SARM1) were compared between wild-type cultures of the two neuron types (fig. 4a). Whereas levels of SARM1 appeared to be similar between the two neuron types (fig. 4b, f), NAMPT (nicotinamide phosphoribosyltransferase), the rate-limiting enzyme in the NAD biosynthetic pathway from Nicotinamide (NAM), was significantly lower in DRG neurons compared to SCG neurons (fig. 4b, d). Furthermore, NMNAT2, the enzyme converting NMN to NAD, showed a non-significant trend towards lower expression in SCGs than DRGs (fig. 4b, e). As a result, the NMNAT2:NAMPT ratio, which is likely to influence levels of the endogenous SARM1 activator NMN, was significantly lower in SCG cultures with an effect size of more than twofold (fig. 4c). The higher ratio in DRG neurons could explain why NMNAT2 levels can be reduced further than in SCG neurons before becoming rate limiting, thereby lowering the likelihood of NMN accumulation and consequent SARM1 activation when NMNAT2 levels are chronically low. This could explain, at least partly, the absence of any NAD(P) depletion and neurite outgrowth defect in DRG neurons from *Nmnat2^gtBay/gtE^* mice.

**Fig. 4.**
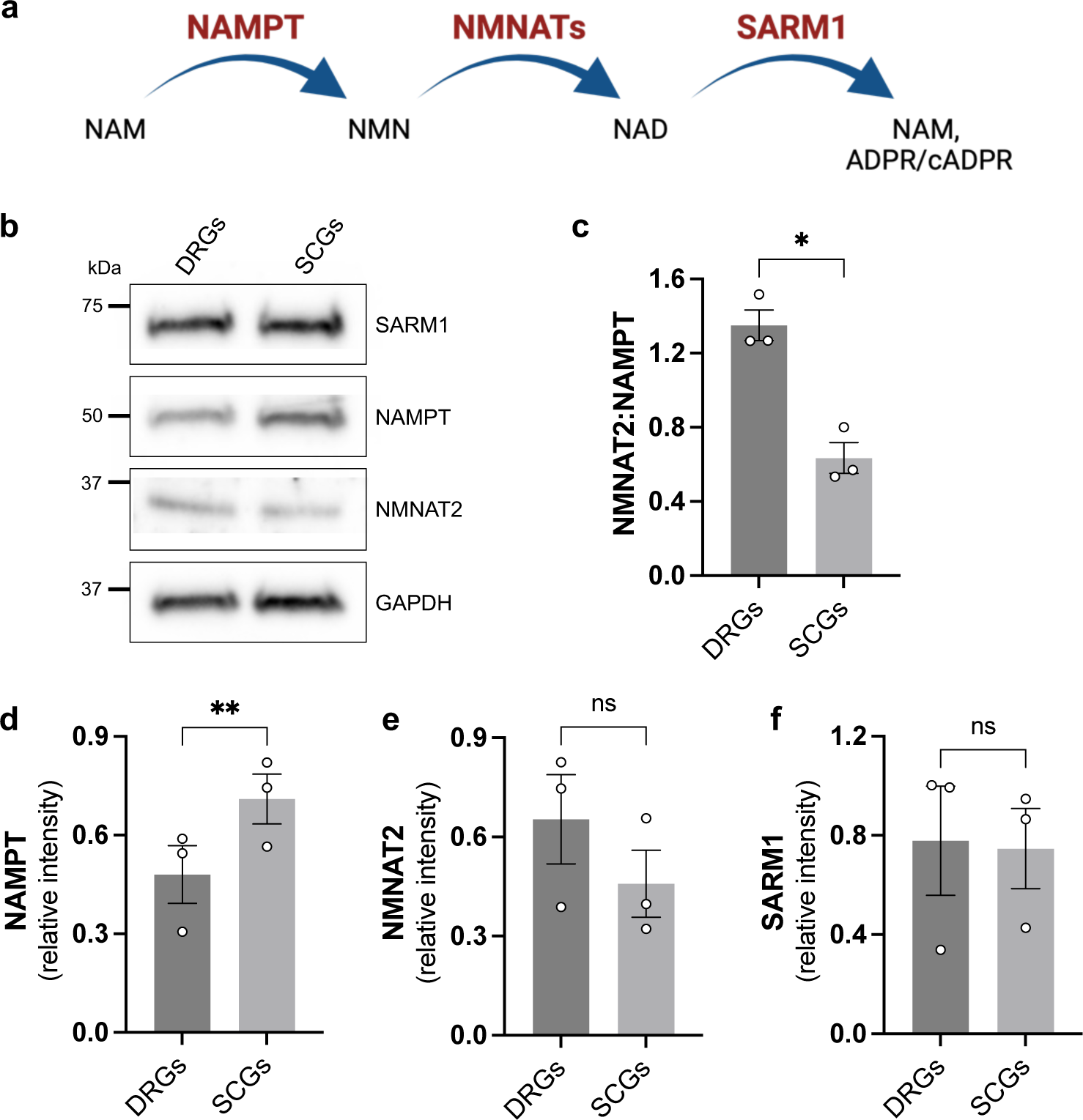
Higher NMNAT2 to NAMPT ratio in DRG vs SCG neurons. (**a**) Pathway of NAD synthesis from precursor NAM and NAD consumption by SARM1. (**b**). Representative immunoblot of wild-type SCG and DRG extracts probed for NAMPT, NMNAT2, SARM1 and GAPDH (loading control). Cultures were collected at DIV7. (**c**) NMNAT2:NAMPT ratio (mean ± SEM; n = 3; *p < 0.05 paired t-test). (**d-f**) Quantification of NAMPT, NMNAT2 and SARM1 levels normalised to GAPDH (mean ± SEM; n = 3; **p < 0.01 and ns (not significant) = p > 0.05, paired t-test).

### NR causes a SARM1-dependent NAD depletion in SCG neurites from Nmnat2^gtBay/gtE^ mice

The data provided thus far support the hypothesis that SARM1 can be chronically activated without causing degeneration, at least in shorter axons. We next sought to investigate whether the balance could be further shifted in favour of SARM1 activation by elevating levels of its endogenous activator, NMN, when there is insufficient NMNAT2 to convert all of it rapidly to NAD.

First, wild-type primary DRG cultures were supplemented with the NAD precursors NR and NAM in order to identify conditions that lead to NAD accumulation, thereby reflecting a prior increase in NMN. NAM is converted to NMN by the enzyme NAMPT, while NR is converted to NMN by nicotinamide riboside kinase 1/2 (NRK1 and NRK2). While NR as well as the combination of NR and NAM resulted in a marked increase in NAD(P) levels after a 24-hour treatment, NAM application alone did not significantly increase the levels of either metabolite (supplementary fig. 2). This observation could be due to NAMPT already operating at saturation, as it is the rate-limiting enzyme for NAD biosynthesis in mammals and NAM is already present in the culture medium, so any further increase in NAM does not increase the rate of NAD production. For this reason, NR alone was used in subsequent experiments (fig. 5a).

**Fig. 5.**
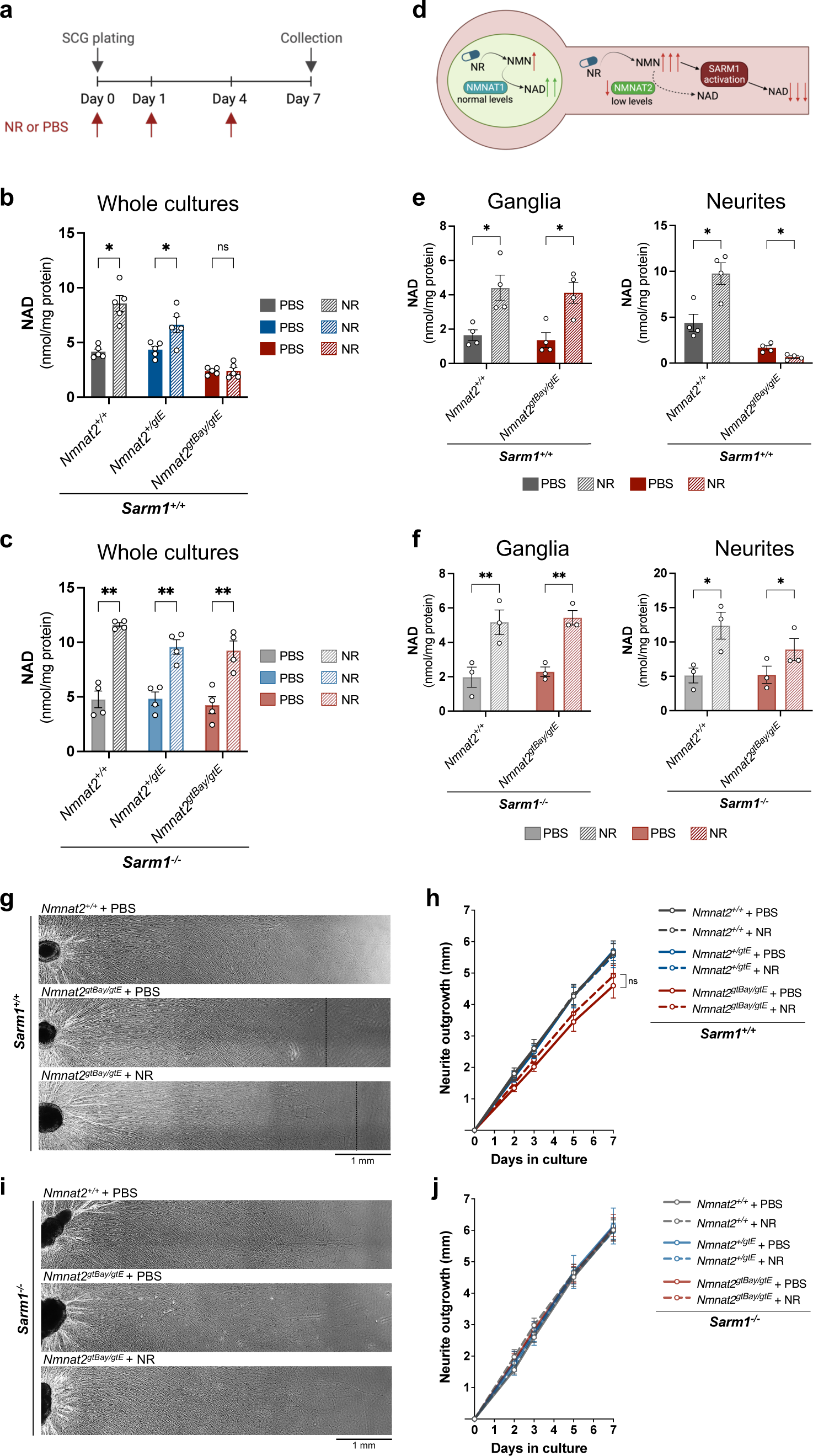
NR causes SARM1-dependent NAD depletion in SCG neurites from Nmnat2^gtBay/gtE^ mice. (**a**) Timeline of NR (2 mM) or PBS administration and collection of SCG cultures. (**b**) NAD levels in whole SCG explants of the indicated genotypes on a *Sarm1^+/+^* background (mean ± SEM; n = 5; *p < 0.05 and ns (not significant) = p > 0.05, multiple paired t-tests for PBS vs NR with Holm-Šídák correction method). (**c**) NAD levels in whole SCG explants of the indicated genotypes on a *Sarm1^-/-^* background (mean ± SEM; n = 4; **p < 0.01, multiple paired t-tests for PBS vs NR with Holm-Šídák correction method). (**d**) Schematic of NMNAT1 and NMNAT2 localisation in neurons and effects of NR administration. (**e**) NAD levels in SCG ganglia and neurites of the indicated genotypes on a *Sarm1^+/+^* background (mean ± SEM; n = 4; *p < 0.05, multiple paired t-tests for PBS vs NR with Holm-Šídák correction method). (**f**) NAD levels in SCG ganglia and neurites of the indicated genotypes on a *Sarm1^-/-^* background (mean ± SEM; n = 3; **p < 0.01 and *p < 0.05, multiple paired t-tests for PBS vs NR with Holm-Šídák correction method). (**g**) Representative images of neurite outgrowth at DIV7 in SCG explant cultures of the indicated genotypes on a *Sarm1^+/+^* background after administration of NR (2 mM) or PBS control. (**h**) Quantification of neurite outgrowth in SCG explant cultures of the indicated genotypes on a *Sarm1^+/+^* background, between DIV0 and DIV7 (mean ± SEM; n = 5; ns (not significant) = p > 0.05, two-way repeated measures ANOVA with Tukey’s multiple comparisons test for between genotype effects at each time point). (**i**) Representative images of neurite outgrowth at DIV7 in SCG explant cultures of the indicated genotypes on a *Sarm1^-/-^* background after administration of NR (2 mM) or PBS control. (**j**) Quantification of neurite outgrowth in SCG explant cultures of the indicated genotypes on a *Sarm1^-/-^* background, between DIV0 and DIV7 (mean ± SEM; n = 3; ns (not significant) = p > 0.05, two-way repeated measures ANOVA with Tukey’s multiple comparisons for between genotype effects at each time point).

Supplementation with NR also significantly increased NAD levels in wild-type and *Nmnat2^+/gtE^* whole SCG cultures but had no effect in *Nmnat2^gtBay/gtE^* whole SCG cultures. NR and PBS-treated *Nmnat2^gtBay/gtE^* cultures were indistinguishable (fig. 5b). The lack of NAD increase in *Nmnat2^gtBay/gtE^* SCG cultures in response to NR could be attributed to low NMNAT2 expression being unable to utilise the excess of NMN for NAD synthesis. However, absence of SARM1 restored the ability of *Nmnat2^gtBay/gtE^* neurons to increase NAD levels following NR treatment (fig. 5c). This suggests that the lack of NAD accumulation in the first instance is not due to lower NAD synthesising capacity, as this is also lower in the *Sarm1*^-/-^ cultures, but rather to the lower capacity to remove NMN, whose accumulation activates SARM1 NADase. Unfortunately, it was not feasible to measure NMN levels in these primary cultures to confirm an increase. The limited number of pups carrying the genotypes of interest, coupled with the reduced viability of *Nmnat2^gtBay/gtE^* animals, hindered the acquisition of sufficient material necessary for metabolite analysis.

The observation that NR administration had no net effect on NAD levels in *Nmnat2^gtBay/gtE^* cultures led to the hypothesis that the response to NR could differ between neurites and cell bodies, as cell bodies also have NMNAT1 to convert NMN to NAD (fig. 5d). For this reason, separate metabolite measurements were made in cell body and neurite compartments following NR administration in wild-type and *Nmnat2^gtBay/gtE^* SCG cultures. Interestingly, NR administration caused NAD to decline significantly in neurites with sub-heterozygous NMNAT2 expression (fig. 5e). In contrast, the cell bodies of *Nmnat2^gtBay/gtE^* SCG cultures showed increased NAD when supplemented with NR, consistent with conversion of NMN to NAD by NMNAT1, the nuclear NMNAT isoform (fig. 5e). Given that NMNAT1 levels are unaffected in *Nmnat2^gtBay/gtE^* cultures, this isoform is able to utilise the NMN derived from NR, resulting in the increased levels of NAD seen in the cell bodies (fig. 5d). Hence, the observation that NAD levels in whole SCG *Nmnat2^gtBay/gtE^* cultures do not change with NR supplementation is likely to reflect the net effect of both increasing NAD in cell bodies and decreasing it in neurites. In *Nmnat2^gtBay/gtE^* neurites lacking SARM1, NAD levels were increased following NR administration (fig. 5f). A similar trend was observed with NADP, although the changes were less marked (supplementary fig. 3). Collectively, these observations strongly suggest that under conditions of inadequate NMNAT2, the accumulated NMN resulting from NR administration further activates SARM1, leading to a depletion of NAD levels in *Nmnat2^gtBay/gtE^* neurites.

Interestingly, despite NR leading to further activation of SARM1 in neurites with low NMNAT2, no morphological changes were seen in these cultures. There were no signs of frank axon degeneration (data not shown) and somewhat counter-intuitively the neurite outgrowth defect was actually slightly improved, albeit not significantly (fig. 5g, h). We propose that this could be attributed to the timing of NR administration, with transient increases in NAD levels occurring shortly after NR administration giving neurites an initial growth spurt before SARM1 activation and NAD depletion take place. Alternatively, the increase in NAD levels occurring in *Nmnat2^gtBay/gtE^* cell bodies following NR administration could account for the improvement in the outgrowth phenotype. Interestingly, increased levels of NAD after NR supplementation had no effect on neurite outgrowth in *Nmnat2^+/+^* and *Nmnat2^+/gtE^* cultures on a *Sarm1*^+/+^ background, or in any *Nmnat2* genotype lacking SARM1 (fig 5h, i). This indicates that enhanced NAD alone is not sufficient to boost neurite outgrowth.

Despite DRG cultures from *Nmnat2^gtBay/gtE^* mice having no phenotype in terms of baseline NAD levels, NR administration had similar effects to those seen in the more susceptible SCG cultures, causing NAD to decline in neurites with sub-heterozygous NMNAT2 expression (supplementary fig. 4). This suggests that in DRG neurons 30% of the C57BL/6 NMNAT2 expression level is insufficient to activate SARM1 and cause NAD(P) depletion under basal conditions. However, boosting NMN levels with the precursor NR, is able to tip the balance further in favour of SARM1 activation and lower NAD, specifically in the neurites, where NMNAT2 is limiting.

## Discussion

The findings presented here indicate that chronically low NMNAT2 expression causes sub-lethal SARM1 activation, which can be enhanced by the NMN precursor NR. SARM1 activation is thus not a binary, all or nothing response but appears to lie on a spectrum, where partial activation does not translate to frank axon degeneration but is still detectable at the molecular level. While SARM1 activation seems to occur when NMNAT2 levels fall below a heterozygous threshold of 50% of normal C57BL/6 mouse expression, it is unknown how this level compares to the human spectrum of NMNAT2 expression. It is also possible that a less pronounced decrease from mean NMNAT2 levels or activity has comparable outcomes in humans, especially considering that human axons are longer, and exposed to multiple neurodegenerative stresses over a substantially longer lifespan.

Our work supports the hypothesis that low NMNAT2 levels compromise prenatal survival in a SARM1-dependent manner, demonstrating that targeting SARM1 can be beneficial not only in conditions of complete [4] but also of partial NMNAT2 loss. This observation reinforces the trend of reduced viability we previously reported [21], potentially strengthened by genetic selection and/or environmental differences following a move of our mice to a new animal facility between the two studies. These findings support the idea that decreased NMNAT2 expression could be more problematic in some people than others, given the widespread variability in genotype and environment within the human population.

Based on the Genome Aggregation Database (gnomAD), the probability of LOF intolerance (pLI) for human NMNAT2 is 0.98 suggesting intolerance of, and selective pressure against hemizygosity. Nevertheless, all six of the parents of the biallelic cases so far reported are neurologically healthy [11–13] suggesting that other genetic and/or environmental factors may modify the outcome to explain their healthy survival despite this selective pressure. This appears to parallel the extreme variability between outcomes in *Nmnat2^gtBay/gtE^* mice (albeit at sub-heterozygous level), where some pups are non-viable while the ones born alive remain overtly normal throughout life. Interestingly, this is the case despite our mice having greater genetic and environmental homogeneity than that in the human population.

Our study has also shown that the NAD precursor supplement NR lowers NAD instead of increasing it in neurites expressing sub-heterozygous levels of NMNAT2. While NR supplementation is likely to be harmless and potentially beneficial in the majority of the population, (with no toxicity being reported so far in human studies [30]), our findings suggest that NR and possibly other NAD precursors could be problematic in a subset. As well as the known human cases with mutations in *NMNAT2*, there are other conditions where the levels, activity, or transport of the protein could be compromised. The widespread variability of human *Nmnat2* expression [15] raises the possibility that individuals at the lower end of the spectrum could be at increased risk of SARM1 activation and potential neurotoxicity as a result of NR supplementation.

In addition, ageing has been shown to decrease axonal transport of NMNAT2, at least in mice [31], while inhibition of protein synthesis [1] and mitochondrial dysfunction [22,32] decrease NMNAT2 levels in axons. Thus, the elderly and individuals undergoing treatments that are likely to inhibit axonal transport, including some chemotherapy treatments [33], or people with other axonal transport deficiencies, such as mutations in genes encoding motor proteins or tubulin cofactors [34], could be among the risk groups. Despite the SARM1-dependent NAD loss in *Nmnat2^gtBay/gtE^* neurites following NR administration, no effects on neurite morphology were observed. However, these findings raise important questions as to whether prolonged SARM1 activation and NAD depletion within axons, resulting from NR supplementation, could cause axon degeneration in humans, where axons are much longer, are exposed to many more stressful stimuli, and need to be retained for many decades. In support of this, we previously showed that neurites with sub-heterozygous NMNAT2 expression are more susceptible to other neurodegenerative stresses [21,22].

Furthermore, the data presented here highlight the need to examine the long-term implications of sub-lethal SARM1 activation. SARM1 activation in the absence of axon death has been reported previously in acute models, evident through accumulation of the SARM1 marker cADPR following administration of NMN analogue CZ-48 [5,29], low doses of mitochondrial toxins [35] and treatment with NR in neurons overexpressing the enzyme NRK1 [6] and our data now indicate that partial SARM1 activation can also occur chronically. Intact NMNAT activity is able to compensate for increased SARM1-dependent NAD consumption, however prolonged SARM1 activation and accumulation of associated products, such as cADPR and NAADP (nicotinic acid adenine dinucleotide phosphate), could affect cellular physiology leading to functional defects despite the lack of morphologically visible neurotoxicity. When NMNAT activity is compromised, as we describe here, prolonged SARM1 activation and NAD depletion are expected to have more severe consequences. For instance, chronic SARM1 activation likely results, or at least contributes, to the behavioural phenotypes and deficits in peripheral axon numbers previously reported in the *Nmnat2^gtBay/gtE^* mice [21]. In support, a SARM1-dependent increase in cADPR levels has been reported in sciatic nerves of 2-month-old mice harbouring the human *NMNAT2* LOF mutations (*Nmnat2^V98M/R232Q^*), with SARM1 being required for the neuropathy phenotypes in these mice [13]. Thus, both reduced levels and activity of NMNAT2 can chronically activate SARM1 and compromise neuronal health and survival. It is likely that this will be more pronounced in longer-lived human axons and can potentially worsen with age. Finally, assessing the levels of relevant metabolites in accessible tissues such as blood or CSF could serve as the basis of screening tools for conditions involving SARM1 activation in humans.

The present study has provided evidence in support of chronic, sub-lethal SARM1 activation in non-degenerating axons. What sub-lethal SARM1 activation means in structural terms is nonetheless unknown. One possibility is that fewer SARM1 octamers exist in an active conformation compared to a fully activated state that leads to degeneration. Alternatively, the break of the ARM-TIR lock, which mediates the conformational change required for SARM1 activation might itself be partial. Another possibility would be transient activation when NMN binds followed by deactivation when it dissociates, or is replaced by NAD. Structural studies will thus be instrumental in answering this question. Moreover, the absence of degeneration could be attributed to compensatory mechanisms arising in response to constitutively low NMNAT2 expression. In support, we previously showed that acute depletion of a single *Nmnat2* allele causes axon degeneration in primary cultures [3], whereas constitutively lower levels of NMNAT2 are compatible with axon survival.

In summary, we have shown that constitutively low NMNAT2 levels reduce viability in mice and lead to sub-lethal SARM1 activation in morphologically intact axons, characterised by NAD(P) depletion and the development of shorter neurites. The effect of chronic NMNAT2 depletion is not uniform across different neuron types and variations in the NMNAT2 to NAMPT ratio might account for the differential susceptibility. Finally, we argue that supplementation with NR and other NAD precursors may need additional safety studies in conditions of reduced NMNAT function. Importantly, compromised or reduced NMNAT2 activity could have a more profound effect in human axons considering their greater length and longer lifespan. Although LOF mutations in the *NMNAT2* gene are rare, the widespread variability of *NMNAT2* mRNA expression reported in humans, together with the multitude of pathological and physiological situations that can compromise NMNAT2 transport or synthesis, could mean that the effects of NMNAT2 on SARM1 activation might be more widespread than previously anticipated. Finally, the findings of this study raise important questions as to what other neurodegenerative stresses can partially activate SARM1, to what extent they contribute to sporadic neurodegenerative diseases and importantly, if early detection and intervention is possible in humans.

## Acknowledgements

We thank Astra Zeneca for synthesising and providing PC6 and Yi-Ping Hsueh for providing the SARM1 monoclonal antibody.

## Author contribution

Christina Antoniou: conceptualisation, data acquisition, data analysis, data interpretation, study design, writing—original draft, writing—review and editing. Andrea Loreto: data acquisition, data interpretation, study design, writing—review and editing. Jonathan Gilley: data interpretation, study design, writing—review and editing. Elisa Merlini: data acquisition, writing—review and editing. Giuseppe Orsomando: data interpretation, writing—review and editing. Michael P Coleman: conceptualisation, data interpretation, study design, supervision, writing—original draft, writing—review and editing. All authors read and approved the final manuscript.

## Funding

C.A. is funded by the MRC DTP Studentship and Gates Foundation; A.L. is funded by the Wellcome Trust [Grant number 210904/Z/18/Z]; E.M. is funded by the Cambridge Trust; J.G. is funded by ALS Finding a Cure and the ALS Association 959996; G.O. is funded by the Italian Grants RSA 2020-2022 from UNIVPM. M.P.C. is funded by the John and Lucille van Geest Foundation.

## Data Availability

The datasets generated and analysed during the current study are available from the corresponding author upon reasonable request.

## Declarations

### Competing Interests

MPC consults for Nura Bio and Drishti Discoveries and the Coleman group is part funded by AstraZeneca for academic research projects but none of these activities relate to the study reported here.

### Ethics Approval

Animal work was approved by the University of Cambridge and performed in accordance with the Home Office Animal Scientific Procedures Act (ASPA), 1986 under project licence P98A03BF9.

### Consent to Participate

Not applicable.

### Consent for Publication

Not applicable.

