## Supplemental files for "Chronically low NMNAT2 expression causes sub-lethal SARM1 activation and altered response to nicotinamide riboside in axons"

Molecular Neurobiology

Christina Antoniou[1], Andrea Loreto[1], Jonathan Gilley1, Elisa Merlini[1], Giuseppe Orsomando[2] and Michael P Coleman[1]

**Author affiliations:**

1. John van Geest Centre for Brain Repair, Department of Clinical Neurosciences, University of Cambridge, Forvie Site, Robinson Way, CB2 0PY Cambridge, UK

2. Department of Clinical Sciences (DISCO), Section of Biochemistry, Polytechnic University of Marche, Via Ranieri 67, Ancona 60131, Italy

**Live births**

| x | Genotype | Viable Offspring: Observed | Viable Offspring: Expected |
|---|---|---|---|
| | *Nmnat2$^{+/+}$;Sarm1$^{+/+}$* | **561** | 558.5 |
| *Nmnat2$^{+/gtBay}$;Sarm1$^{+/+}$* | *Nmnat2$^{+/gtBay}$;Sarm1$^{+/+}$* | **572** | 558.5 |
| x | *Nmnat2$^{+/gtE}$;Sarm1$^{+/+}$* | **605** | 558.5 |
| *Nmnat2$^{+/gtE}$;Sarm1$^{+/+}$* | *Nmnat2$^{gtBay/gtE}$;Sarm1$^{+/+}$* | **496** | 558.5 |

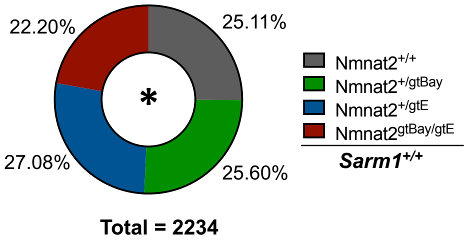


**Supplementary figure 1** Sub-heterozygous NMNAT2 expression in mice reduces the number of live births. Genotype frequencies of viable offspring from crosses between *Nmnat2$^{+/gtE}$* and *Nmnat2$^{+/gtBay}$* mice, on a *Sarm1$^{+/+}$* background. The observed birth frequencies are significantly different from expected frequencies: $\chi^2$ = 11.203, d.f. = 3, p = 0.0107. Viable offspring numbers include animals between P0-P3 and post-weaning, combining the live births recorded in our previous study (before the transfer to the new animal facility) and the current study (after the transfer to the new animal facility).

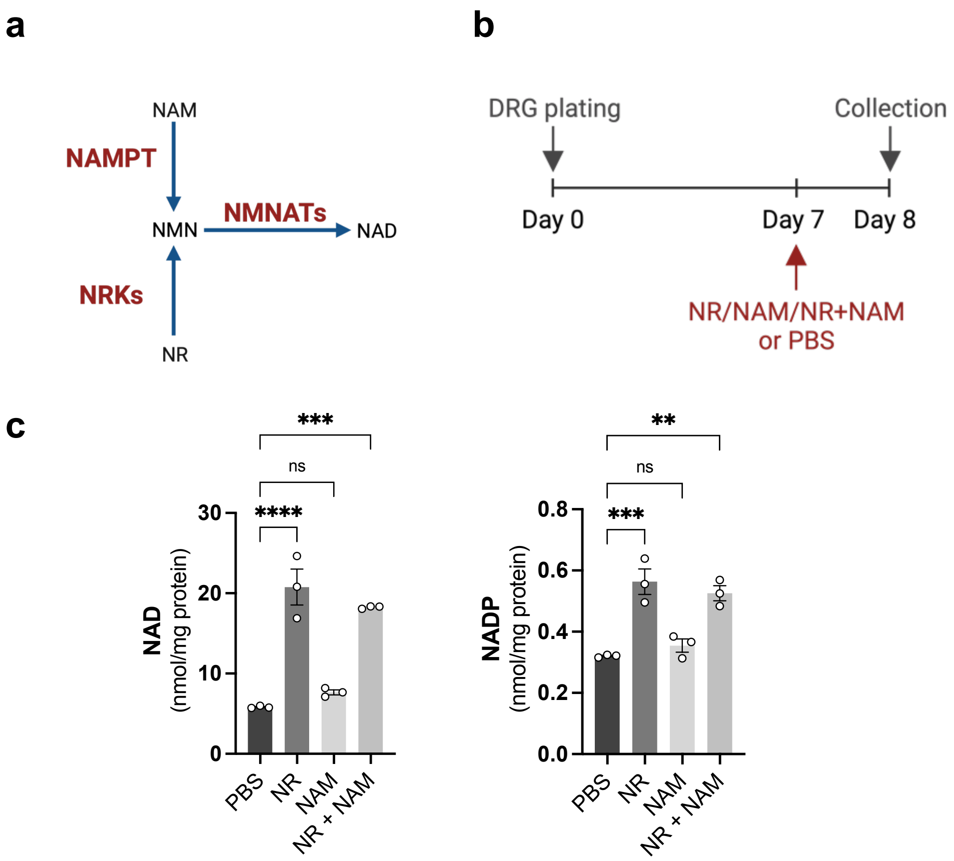


**Supplementary figure 2** Testing supplementation of NAD precursors in primary neuronal cultures. (**a**) Pathway of NAD synthesis from precursors NAM (nicotinamide) and NR (nicotinamide riboside). (**b**) Timeline of NAD precursor administration and collection of whole DRG cultures. (**c**) NAD and NADP levels in wild-type DRG explants following administration of NR (2 mM), NAM (1 mM), combination of NR and NAM or PBS control (mean ± SEM; n = 3; ****$p < 0.0001$, ***$p < 0.001$, **$p < 0.01$ and ns (not significant) = $p > 0.05$, one-way ANOVA with Dunnett's test comparing means to PBS control).

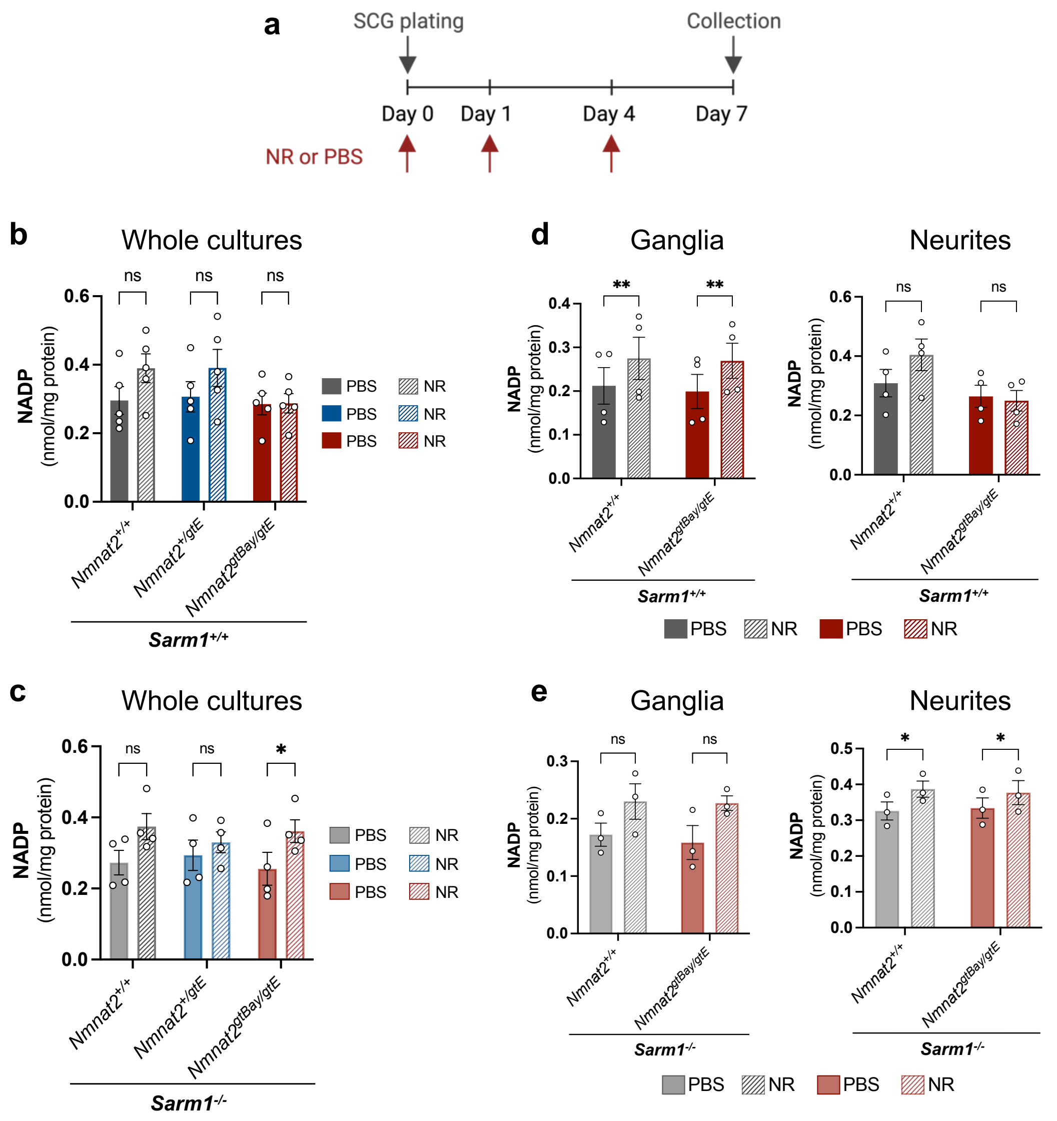


**Supplementary figure 3** NR administration does not increase NADP levels in DRG whole cultures and neurites from *Nmnat2$^{gtBay/gtE}$* mice. (**a**) Timeline of NR (2 mM) or PBS administration and collection of SCG cultures. (**b**) NADP levels in SCG explants of the indicated genotypes, all on a *Sarm1$^{+/+}$* background (mean ± SEM; n = 5; ns (not significant) = p > 0.05, multiple paired t-tests for PBS vs NR with Holm-Šídák correction method). (**c**) NADP levels in whole SCG explants of the indicated genotypes, all on a *Sarm1$^{-/-}$* background (mean ± SEM; n = 4; *p < 0.05 and ns (not significant) = p > 0.05 multiple paired t-tests for PBS vs NR with Holm-Šídák correction method). (**d**) NADP levels in SCG ganglia and neurites of the indicated genotypes, all on a *Sarm1$^{+/+}$* background (mean ± SEM; n = 4; **p < 0.01 and ns (not significant) = p > 0.05, multiple paired t-tests for PBS vs NR with Holm-Šídák correction method). (**e**) NADP levels in SCG ganglia and neurites of the indicated genotypes, all on a *Sarm1$^{-/-}$* background (mean ± SEM; n = 3; *p < 0.05 and ns (not significant) = p > 0.05, multiple paired t-tests for PBS vs NR with Holm-Šídák correction method).

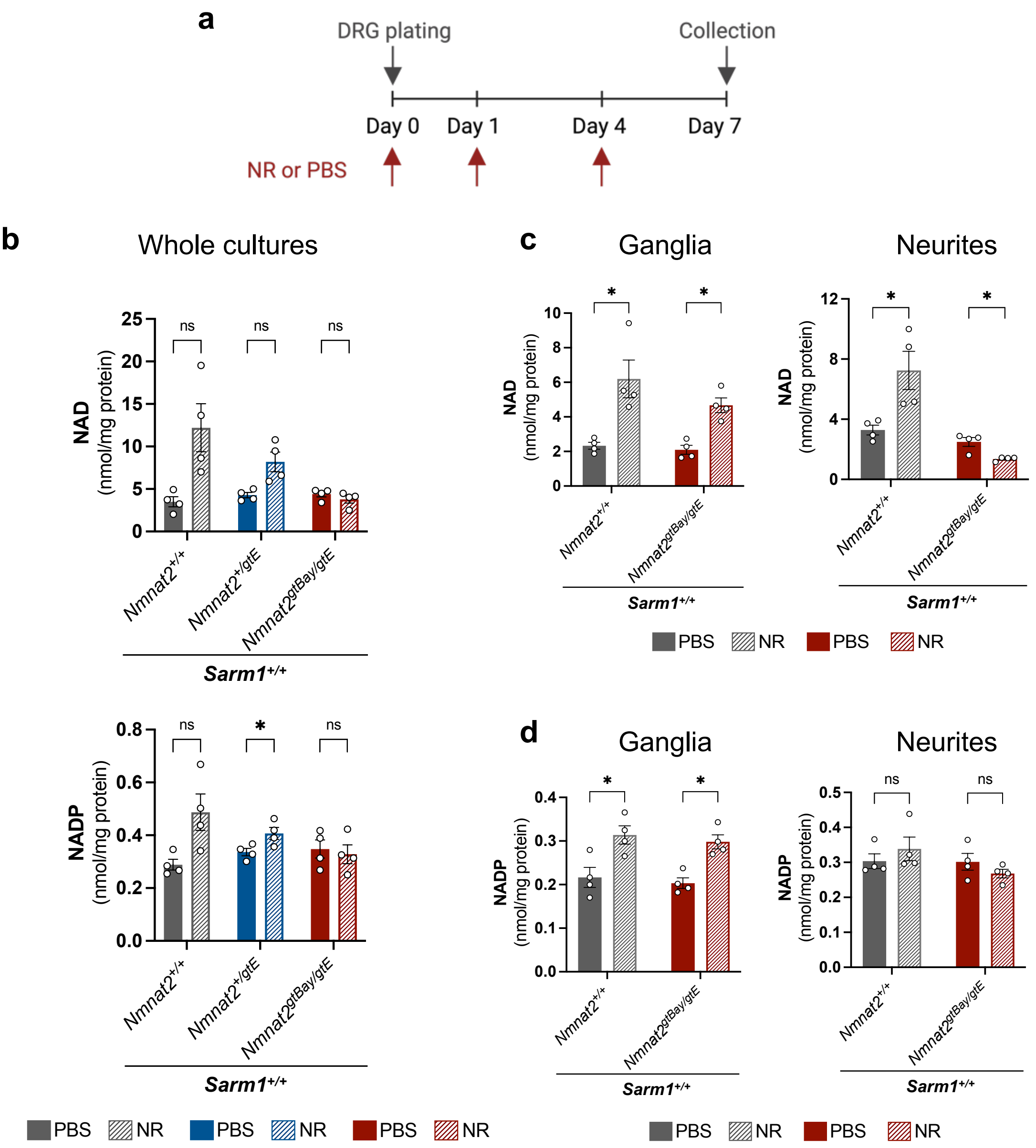


**Supplementary figure 4** NR causes a SARM1-dependent NAD depletion in DRG neurites from Nmnat2$^{gtBay/gtE}$ mice. (a) Timeline of NR (2 mM) or PBS administration and collection of DRG cultures. (b) NAD and NADP levels in whole DRG explants of the indicated genotypes, all on a *Sarm1*$^{+/+}$ background (mean ± SEM; n = 4; *p < 0.05 and ns (not significant) = p > 0.05, multiple paired t-tests for PBS vs NR with Holm-Šídák correction method). (**c**) NAD levels in DRG ganglia and neurites of the indicated genotypes all on a *Sarm1*$^{+/+}$ background (mean ± SEM; n = 4; *p < 0.05 multiple paired t-tests for PBS vs NR with Holm-Šídák correction method). (**d**) NADP levels in DRG ganglia and neurites of the indicated genotypes, all on a *Sarm1*$^{+/+}$ background (mean ± SEM; n = 4; *p < 0.05 and ns (not significant) = p > 0.05, multiple paired t-tests for PBS vs NR with Holm-Šídák correction method).

### Uncropped immunoblot images for figure 2d

Membrane probed for SARM1

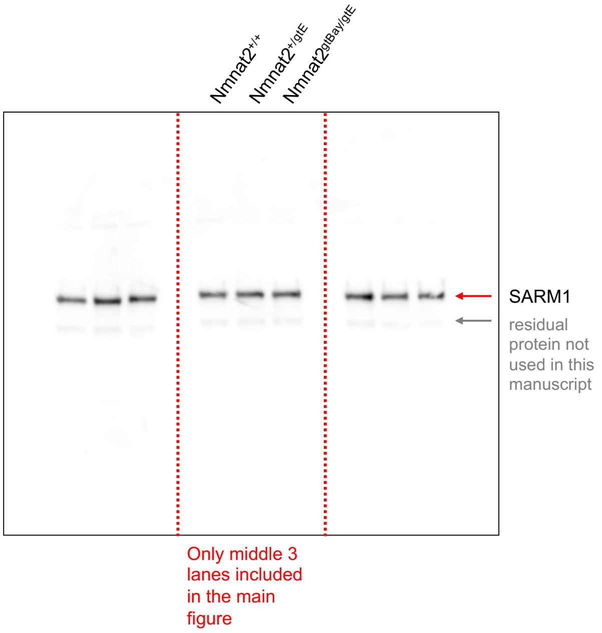


Membrane probed for GAPDH

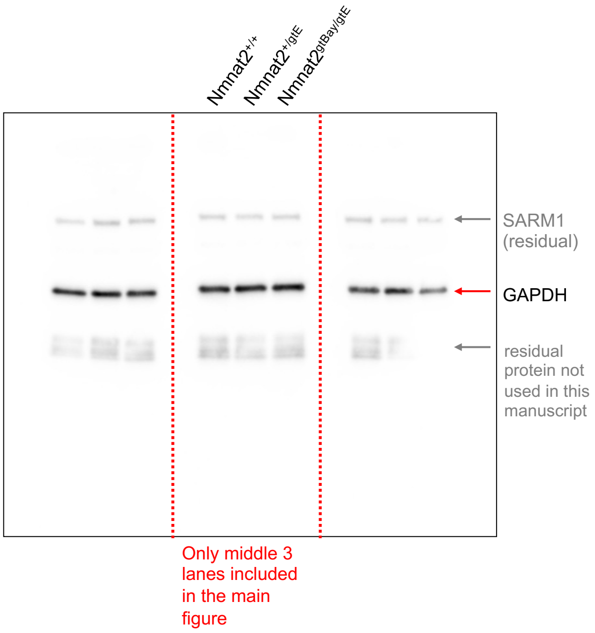


### Uncropped immunoblot images for figure 4b

Membrane probed simultaneously for SARM1 and NAMPT

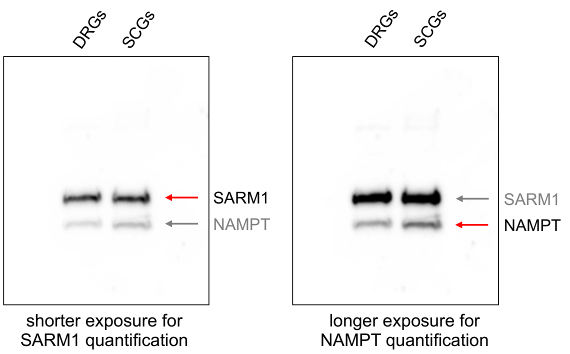


Membrane probed for NMNAT2

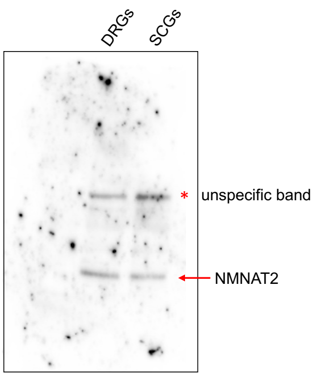


Membrane probed for GAPDH

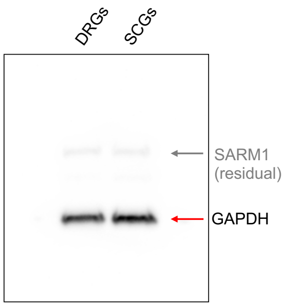
